# Behavioural and neural interactions between objective and subjective performance in a Matching Pennies game

**DOI:** 10.1101/598466

**Authors:** Benjamin James Dyson, Cecile Musgrave, Cameron Rowe, Rayman Sandhur

## Abstract

To examine the behavioural and neural interactions between objective and subjective performance during competitive decision-making, participants completed a Matching Pennies game where win-rates were fixed within three conditions (*win > lose, win = lose, win < lose*) and outcomes were predicted at each trial. Using random behaviour as the hallmark of optimal performance, we observed item (*heads*), contingency (*win-stay, lose-shift*) and combinatorial (HH, HT, TH, TT) biases across all conditions. Higher-quality behaviour represented by a reduction in combinatorial bias was observed during high win-rate exposure. In contrast, over-optimism biases were observed only in conditions where win rates were equal to, or less than, loss rates. At a group level, a neural measure of outcome evaluation (feedback-related negativity; FRN) indexed the binary distinction between positive and negative outcome. At an individual level, increased belief in successful performance accentuated FRN amplitude differences between wins and losses. Taken together, the data suggest that objective experiences of, or, subjective beliefs in, the predominance of positive outcomes are mutual attempts to self-regulate performance during competition. In this way, increased exposure to positive outcomes (real or imagined) help to weight the output of the more diligent and analytic System 2, relative to the impulsive and intuitive System 1.

---

Competitive decision-making represents a critical test scenario where behavioural biases that have enabled species survival also make an organism more predictable and hence prone to exploitation. These tensions are summarized in the dynamic weighting of two forms of processing systems within the brain: System 1 reflecting relatively impulsive, intuitive and inflexible processing, and System 2 reflecting relatively diligent, analytic and flexible processing (Sloman, 1996; Kahneman, 2011; Williams, Kappen, Hassall, Wright & Krigolson, in presss). Successful behaviour relies heavily on the output of System 2 but the characteristics of the competitive environment may encourage System 1 output instead (Forder & Dyson, 2016). One of the benefits in examining competitive decision-making using simple zero-sum games is that optimal performance predicted by System 2 output can be clearly defined. For example, in games such as Rock, Paper, Scissors (RPS; e.g., Cook, Bird, Lünser, Huck & Heyes, 2012; Dyson, Wilbiks, Sandhu, Papanicolaou & Lintag, 2016) or Matching Pennies (MP; e.g., Belot, Crawford & Heyes, 2013; Aczel, Kekees, Bago, Szollosi & Foldes, 2015) the only way to avoid exploitation with certainty is to play in accordance with a mixed-equilibrium strategy (MES). Here, all responses are distributed equally through the duration of the game and no contingencies exist between one trial and the next (Baek et al., 2013; Bi & Zhou, 2014; Lee, McGreevy & Barraclough, 2005). However it is a common observation that humans find it difficult to behave randomly (e.g., Griessinger & Coricelli, 2015; Heuer, Janczyk, & Kunde, 2010; Terhune & Brugger, 2011) because of exposure to positively auto-correlated or ‘clumpy’ sequences in the natural world (Scheibehenne, Wilke, & Todd, 2011). A second reason why the empirical data fails to demonstrate random behaviour may due to the task itself, in that commonly-used ‘production’ tasks where participants are explicitly asked to behave randomly are beset by logical and methodological problems (see Rapoport & Budescu, 1992, for a discussion). Instead, tasks within which randomness is a means to another end-such as avoiding losses within a competitive game-are proposed as more liberal tests of the ability to express randomness (Rapoport & Budescu, 1992). Moreover, competitive zero-sum games such as RPS and MP also provide a multi-nominal distribution of responses and so it becomes easier to test the degree to which participants are performing in accordance with the optimal conditional probabilities dictated by MES (i.e., for items: 33.3% in RPS, 50% in MP).

In terms of assessing the degree to which individual deviate from random (optimal) performance, a number of metrics are available including the evaluation of item biases, contingency biases, and, combinatorial biases. Item biases simply reflect the over-play of certain responses relative to others, according to the optimal conditional probabilities outlined above. For example, rock enjoys a slight preference in RPS selection (Baek et al., 2013; Dyson et al., 2016; Forder & Dyson, 2016; Wang, Xu & Zhou, 2014), whereas the probability of choosing heads in a coin toss is between 70% - 80% (Bakan, 1940, Goodfellow, 1940, cited in Bar-Hillel, Peer & Acquisti, 2014), implicating a similar item bias in MP. In both cases, the parsimonious explanation for why these particular responses are preferred may simply be the result of a reachability bias or primary effect (Bar-Hillel et al., 2014), where participants are more likely to select the first item within a list of choices. With respect to contingency biases, the outcome of the previous trial *n-1* can influence the nature of response of the subsequent trial *n*, and specifically, the operant conditioning (reinforcement learning) tendencies of *win-stay* and *lose-shift* are key (Thorndike, 1911). Recent evidence has suggested that *lose-shift* behaviour is more robust than *win-stay* behaviour, at least in the context of RPS (Dyson et al., 2016; Forder & Dyson, 2016). This is consistent with more predictable (and hence, poorer) performance following negative outcome, aligned with the greater weighting of System 1-style operation following loss. It is similarly consistent with accounts from evolutionary psychology that the consequences of a misapplied rule following a negative outcome are potentially more catastrophic that the consequences of a misapplied rule following a positive outcome: “there is no time to calculate trajectories and forces when one is being chased by a bear” (West & Lebiere, 2001, p. 236). One obvious concern surrounding this interpretation however is the use of three outcomes in RPS, where only one of which is perceived as positive (i.e., *win*). Therefore, the preponderance of *lose-shift* behaviour may simply be due to increased exposure to negative outcomes (i.e., *lose, draw*). We address this concern here by using MP, where the binary outcomes of each trial are either clearly negative or clearly positive. Finally, combinatorial biases look beyond relatively simple outcome-response associations and consider to what extent participants fail to equally distribute the repetition and alternation of responses across increasingly demanding combination sizes (i.e., duplets, triplets and quartets; see also descriptions of *lag1, lag2, lag3* performance; West & Lebiere, 2001). The ability of participants to approximate the equal distribution of all response combinations appears to be limited to the 4 possible duplets (i.e., HH, HT, TH, TT), rather than the 8 possible triplets or 16 possible quartets available in a binary choice game (Budescu & Rapoport, 1992; Rapoport & Budescu, 1992). This is entirely consistent with the increased working memory demands associated with keeping track of the frequency of individual combinations, as the combination size increases.

If deviations from randomness and sub-optimal System 1-style behaviour are in part predicted by the experience of negative relative to positive outcomes (Dyson et al., 2016; Laakasuo et al., 2015; Mitzenmacher & Upfal, 2005), then a critical component in the expression of higher- or poorer-quality behaviour becomes the relationship between actual and perceived performance. Here, a final form of bias is that participants exhibit over-optimism regarding their own performance (Cazé & van der Meer, 2013). Such *optimism biases* (Sharot, Riccardi, Raio & Phelps, 2007) serve a self-regulatory purpose, biasing attention away from negative outcome and drawing attention towards positive outcomes (contra Sun, Bai, Yu, Zhou, Zhang & Shen, 2015). As such, the belief that one is performing better than one actually is might serve to promote the operation of System 2 rather than System 1, depending on the degree to which the number of positive outcomes is over-estimated.

This optimistic bias is also thought to lie at the heart of the operation of an early event-related potential explicitly associated with the evaluation of outcome: feedback-related negativity (FRN). FRN is an event-related potential (ERP) maximal at fronto-central electrode sites and is a reliable index of the positive or negative nature of trial outcome occurring some 200 – 300 ms after the on-set of outcome presentation (Miltner, Brown & Coles, 1997; Holroyd, Hajack & Larsen, 2006; see Hauser et al., 2014, and, Luft, 2014, for reviews). FRN is often indistinguishable between unambiguously negative (i.e., *lose*) and ambiguously negative (i.e., *draw*) trials (e.g., Dyson, Steward, Meneghetti & Forder, 2019; Holroyd, Hajack & Larsen, 2006; Kreussel, Hewig, Krestchmer, Hecht, Coles, & Miltner, 2012), but both *lose* and *draw* trials generate larger FRN amplitude than unambiguously positive trials (i.e., *win*; e.g., Gentsch, Ullsperger & Ullsperger, 2009; Nieuwenhuis, Holyord, Mol & Coles, 2004). Such data suggest that FRN reflects a binary classification process between ‘worse-than-expected’ and ‘better-than-expected’ outcomes (Holroyd et al., 2006; Kujawa, Smith, Luhmann & Hajack, 2013), where the default expectation is a *win*. While it remains an assumption that the response of the FRN rests on a default optimism bias, the degree to which individual variation in over-confidence interacts with neural and behavioural variation during competition has not been directly explored. To this end, we designed a study with the following key characteristics and predictions.

Participants took part in an MP paradigm, where responses regarding coin side selection (*heads, tails*) and outcome prediction (*win, lose*) were collected at every trial. To assess the degree to which behavioural and neural response to outcome were modulated by objective changes in performance, three conditions were run where win-rates were fixed within each condition (A: *win > lose*, B: *win = lose*, C: *win < lose*). If the experience of more losses than wins invokes the less diligent, System 1-style processing, then we expected to see the number of item biases, contingency biases, and combinatorial biases increase as the number of losses increased (C > B > A). The use of fixed win-rates throughout a block of trials has featured in previous electrophysiological work and allows for an examination of objective expectancy effects that may influence FRN amplitude (Holyroyd & Krigolson, 2007; Müller et al., 2005). For example, Hajcak, Holroyd, Moser & Simons (2005, Experiment 2), Cohen, Elger & Ranganath (2007), and, Holroyd, Nieuwenhuis, Yeung & Cohen (2003) all employed conditions where participants reliably experienced 75% wins and 25% losses, and, 25% wins and 75% losses (see Fielding, Fu & Franz [in press] for a similar 80% - 20% differential between conditions). Once again, if the processes associated with loss are closer aligned to System 1 output, then changes in FRN amplitude as a result of outcome percentage should be observed following wins but not following losses (Cohen et al., 2007; Forder & Dyson, 2016).

However, as Oliveria et al. (2007) point out, fixing win rates across conditions does not control for potential variation in *subjective* expectancy effects. FRN amplitude to negative outcome may remain larger than FRN amplitude to positive outcome-irrespective of the probability of wins or losses- due to the subjective expectancies surrounding winning. By asking about outcome predictions on every trial, within conditions with fixed win rates, this allowed us to test a) the conditions under which an optimism bias was apparent, and, b) the degree to which optimism biases scaled according to the actual win rate of the condition. To make sure that such estimates could not be influenced by current *objectiv*e differences in performance, each of the three conditions was designed to contain an early and late period where win-rates stabilized at 50%. If optimism bias could be demonstrated during these periods, then this would be a stronger test of historic success or failure influencing the perception of current, consistently random performance. Moreover, from a neural point-of-view, behavioural estimates of confidence also allowed us to assess the degree to which variation in over- or under-confidence might predict FRN amplitude variation. Specifically, we proposed that the higher the over-estimate of wins, the larger the FRN difference between wins and losses.

Finally, the presentation of outcome (and hence, the recording of FRN) directly following an explicit prediction regarding the success or failure of the trial also allowed us to examine the neural representation of match and mismatch between expected and actual outcomes (Bellebaum & Daum, 2008; Hajack et al., 2007). We expected FRN to be larger when there was a mismatch between expected and actual outcomes (Oliveria et al., 2007), that FRN amplitude to mismatches should be larger for unexpected losses relative to unexpected wins (Holyrod et al., 2003), and, that such effects should also be reflected by the further down-stream P300 ERP where losses would be rare with respect to expectancy (as a result of the presumed optimism bias; Hajack et al., 2005).

## Method

### Participants

34 individuals were initially run in the study, with 3 individuals removed due to excessive artifact in the EEG recording. The final sample consisted of 31 individuals (20 female; mean age = 25.03 years, SD = 7.21; 29 right-handers). Participants were entered into a prize draw to win 1 of 3 £25 vouchers. Informed consent was obtained from all participants before testing, and the experiment was approved by the School of Psychology Research Ethics Committee at the University of Sussex (ER/CM698/1).

### Materials

Static images of either the heads and tails side of a British coin were displayed at approximately 6° × 6°, with participants sat approximately 57 cm away from a 22" Diamond Plus CRT monitor (Mitsubishi, Tokyo, Japan). Stimulus presentation was controlled by Presentation 19 (build 03.31.15) and responses were recorded using a keyboard. BFI-10 (Rammstedt & John, 2007) and ER89 (Block & Kremen, 1996) questionnaires were administered at the end of the study.

### Design and procedure

Participants completed 396 trials of MP, divided into three semi-counterbalanced conditions of 132 trials: A (*win > lose*), B (*win = lose*), C (*lose > win*) where each condition was divided into 22 bins of 6 trials (see Figure 1). Each bin had a controlled win-rate between 1/6 and 5/6 but the order of win- and lose-trials within each bin was randomized across participants. By manipulating each bin win-rates, different trajectories could be established across each condition (Sundvall & Dyson, in preparation. For the first and last 18 trials (3 bins x 6 trials) in all conditions, win-rates were 3/6 (50%). Similarly, the 8^th^ and 9^th^, and, 14^th^ and 15^th^ bins also had 50% win-rates, independently of condition. These two sets of bins thus represented intermediate micro-periods where participants received identical exposure to success and failure, but where performance could be influenced by the broader context of the condition. For the remaining bins, and in sets of 4, the win rate rose (*win > lose*), fell (*lose > win*), or, stayed around 50% (*win = lose*). Consequently, participants experienced an overall win-rate of 64% across Condition A, 50% across Condition B, and, 36% across Condition C. Conditions were either presented in order A, B, C, or, C, B, A.

**Figure 1.**
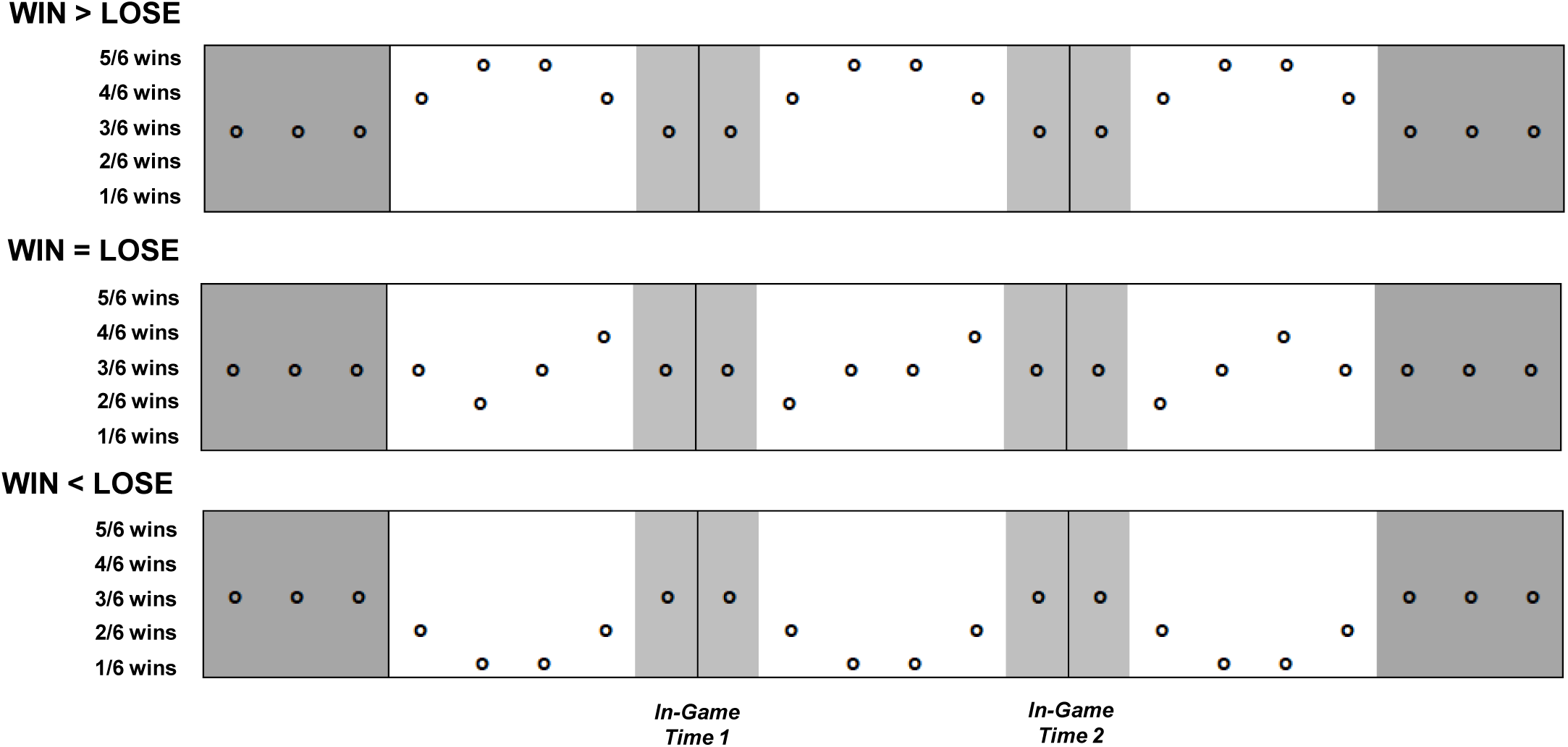
Schematic depicting *win > lose, win = lose*, and, *win < lose* conditions. Each column represents 6 trials within which win rate was fixed between 5/6 and 1/6. The first and last 18 trials were set at 3/6 (50% win rate) for all conditions, as were two sets of 12 trials within the middle of each condition. Modulation of the remaining 54 trials created the differential win-rate experiences.

At each trial, participant score was displayed for 500ms at the bottom right of the screen. Following visual prompts, the participant was then asked to choose a coin side (*heads, tails*) and then to predict the result of the trial (*win, lose*). Their coin side selection was then displayed for 2000ms. This was followed by a clearing of the screen for 500ms followed by visual feedback in the form “WIN” or “LOSE” for 1000ms. At the end of each condition, the participant was asked to predict how many trials they would expect to win if they were to play another 50 trials.

### ERP recording

Electrical brain activity was continuously digitized using a 64 channel ANT Neuro amplifier and a 1000 Hz sampling rate. Horizontal and vertical eye movements were also recorded using channels placed at the outer canthi and at inferior orbits, respectively. Data processing was conducted using BESA 5.3 Research (MEGIS; Gräfelfing, Germany). The contributions of both vertical and horizontal eye movements were reduced from the EEG record using the VEOG and HEOG artefact options in BESA. Following average referencing and using a 0.1 Hz (12 db/oct; zero phase) to 30 Hz (24 db/oct; zero phase) filter, epochs were baseline corrected according to a 200 ms pre-feedback presentation window and neural activity was examined for 800 ms post-feedback presentation. Epochs were rejected on the basis of amplitude difference exceeding 100 μV, gradient between consecutive time points exceeding 75 μV, or, signal lower than 0.01 μV, within any channel. FRN mean amplitudes were calculated on the basis of a 50 ms window centered around the peak latency reported for each specific condition, defined across 225 – 350 ms for nine fronto-central electrodes (F1, Fz, F2, FC1, FCz, FC2, C1, Cz, C2; after Forder & Dyson, 2016; Ma et al, 2014). P3a was measured from the same sites, using a 50 ms mean amplitude window centered around condition peak latencies across 250 – 500 ms.

## Results

### Behavioural Data

As a measure of item bias (see Table 1), the proportion of heads selection was compared across the three conditions (*win > lose, win = lose, win < lose*) via a one-way repeated-measures ANOVA. There was no main effect of condition: F(2,60) = 1.51, MSE =.017, *p* =.229, □_p_^2^ =.048, although was the average proportion of heads selection (52.92%) was significantly different from 50% (*t*[30] = 2.16, *p* =.039), indicating some group item bias. As a measure of contingency bias, the proportion of *win-stay* and *lose-shift* responses were also compared across the three conditions via a two-way repeated-measures ANOVA. There was no main effect of condition: F(2,60) = 0.47, MSE =.010, *p* =.629, □_p_^2^ =.015, no main effect of response: F(1,30) = 0.31, MSE =.013, *p* =.483, □_p_^2^ =.010, and no interaction: F(2,60) = 0.77, MSE =.015, *p* = .469, □_p_^2^ =.025. The average proportion of outcome-strategy bias (53.38%) was however significantly different from 50% (*t*[30] = 2.62, *p* =.014), indicating a slight tendency to deploy *win-stay* / *lose-shift*. As a measure of expressing duplet, triplet and quartet combination bias, standardized ^1^ absolute deviation from expected percentages for each of the three combination types (25.00%, 12.50%, 6.25%, respectively) was calculated in each of three conditions (see Figure 2a; also see Table 2 for a breakdown of proportion by individual combinations). A main effect of combination: F(2,60) = 74.30, MSE <.001, *p* <.001, □_p_^2^ =.712, and interaction between condition x combination were revealed: F(4,120) = 2.76, MSE <.001, *p* =.031, □_p_ ^2^ =.084, in the absence of a main effect of condition: F(2,60) = 0.40, MSE =.004, *p* =.961, □_p_^2^ =.001. Performance became closer to random the smaller the combination analysis size (duplet < triplet < quartet; Tukey’s HSD, *p* <.05), while only duplet performance became closer to random as win rates increased ([win > lose, A] < [win < lose, C]; Tukey’s HSD, p <.05).

**Table 1.** Summary behavioural statistics. Standard error in parenthesis.

|  | Proportion Responses |  |  | Deviation from randomness |  |  |
| --- | --- | --- | --- | --- | --- | --- |
|  | *Heads* | *Win-Stay* | *Lose-Shift* | *Duplet* | *Triplet* | *Quartet* |
| *Win > Lose* | .498 (.029) | .536 (.039) | .512 (.034) | .047 (.007) | .058 (.006) | .068 (.006) |
| *Win = Lose* | .554 (.022) | .569 (.035) | .511 (.028) | .050 (.007) | .056 (.006) | .066 (.006) |
| *Win < Lose* | .535 (.018) | .540 (.037) | .535 (.025) | .052 (.003) | .058 (.004) | .068 (.003) |

|  | Win Rates |  |  |  |  |  |
| --- | --- | --- | --- | --- | --- | --- |
|  | *In-Game Time 1* |  | *In-Game Time 2* |  | *Post-Game Time* |  |
|  | *Actual* | *Predicted* | *Actual* | *Predicted* | *Actual* | *Predicted* |
| *Win > Lose* | .643 | .710 (.042) | .654 | .726 (.045) | .636 | .541 (.038) |
| *Win = Lose* | .500 | .769 (.035) | .500 | .688 (.048) | .500 | .341 (.031) |
| *Win < Lose* | .357 | .640 (.049) | .346 | .616 (.051) | .364 | .263 (.027) |

**Table 2.**
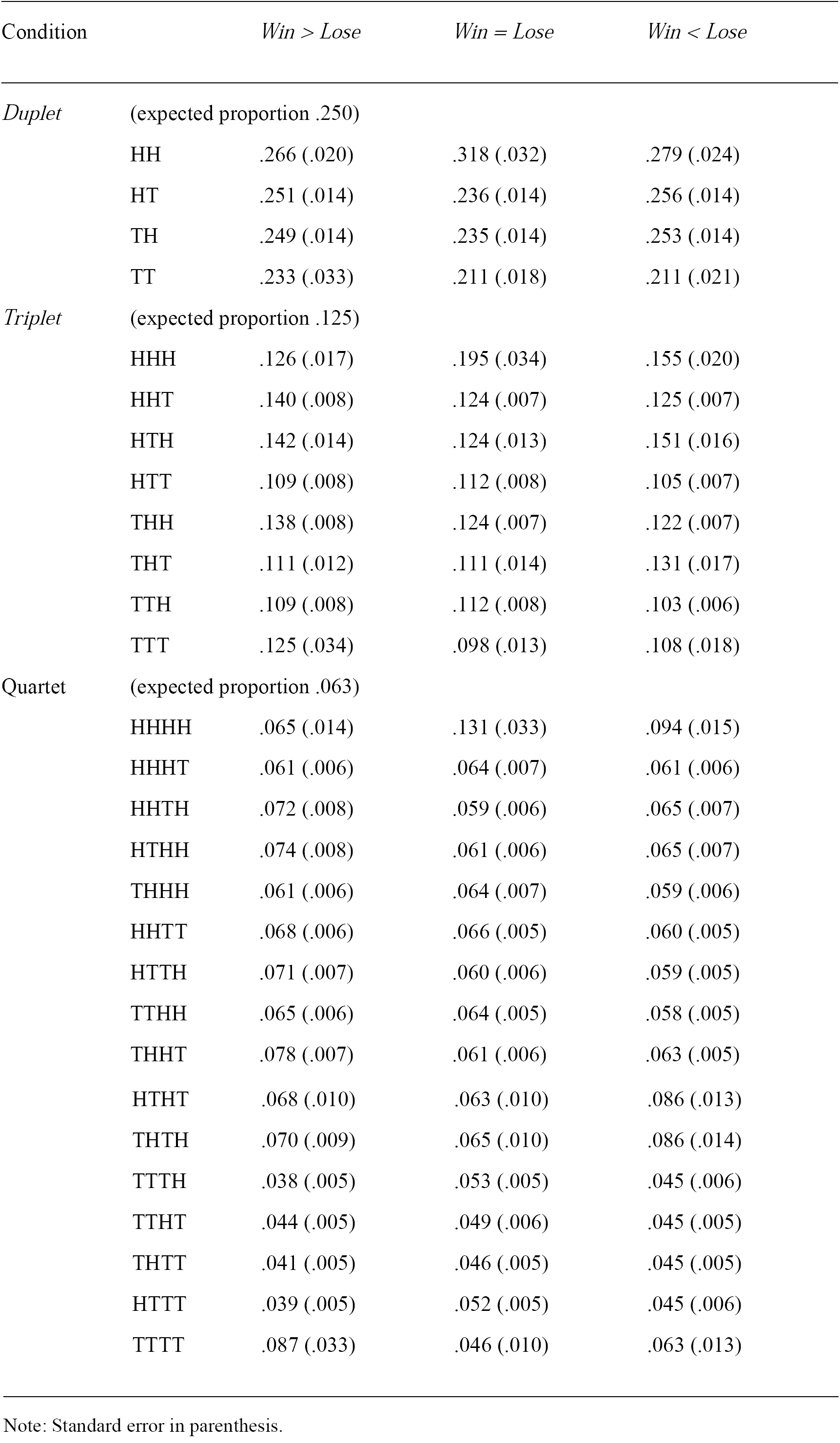
Expected versus observation proportions for all duplet, triplet and quartet combinations

| Condition | | $Win > Lose$ | $Win = Lose$ | $Win < Lose$ |
| --- | --- | --- | --- | --- |
| *Duplet* | (expected proportion .250) |  |  |  |
|  | HH | .266 (.020) | .318 (.032) | .279 (.024) |
|  | HT | .251 (.014) | .236 (.014) | .256 (.014) |
|  | TH | .249 (.014) | .235 (.014) | .253 (.014) |
|  | TT | .233 (.033) | .211 (.018) | .211 (.021) |
| *Triplet* | (expected proportion .125) |  |  |  |
|  | HHH | .126 (.017) | .195 (.034) | .155 (.020) |
|  | HHT | .140 (.008) | .124 (.007) | .125 (.007) |
|  | HTH | .142 (.014) | .124 (.013) | .151 (.016) |
|  | HTT | .109 (.008) | .112 (.008) | .105 (.007) |
|  | THH | .138 (.008) | .124 (.007) | .122 (.007) |
|  | THT | .111 (.012) | .111 (.014) | .131 (.017) |
|  | TTH | .109 (.008) | .112 (.008) | .103 (.006) |
|  | TTT | .125 (.034) | .098 (.013) | .108 (.018) |
| *Quartet* | (expected proportion .063) |  |  |  |
|  | HHHH | .065 (.014) | .131 (.033) | .094 (.015) |
|  | HHHT | .061 (.006) | .064 (.007) | .061 (.006) |
|  | HHTH | .072 (.008) | .059 (.006) | .065 (.007) |
|  | HTHH | .074 (.008) | .061 (.006) | .065 (.007) |
|  | THHH | .061 (.006) | .064 (.007) | .059 (.006) |
|  | HHTT | .068 (.006) | .066 (.005) | .060 (.005) |
|  | HTTH | .071 (.007) | .060 (.006) | .059 (.005) |
|  | TTHH | .065 (.006) | .064 (.005) | .058 (.005) |
|  | THHT | .078 (.007) | .061 (.006) | .063 (.005) |

|  |  |  |  |
| --- | --- | --- | --- |
| HTHT | .068 (.010) | .063 (.010) | .086 (.013) |
| THTH | .070 (.009) | .065 (.010) | .086 (.014) |
| TTTH | .038 (.005) | .053 (.005) | .045 (.006) |
| TTHT | .044 (.005) | .049 (.006) | .045 (.005) |
| THTT | .041 (.005) | .046 (.005) | .045 (.005) |
| HTTT | .039 (.005) | .052 (.005) | .045 (.006) |
| TTTT | .087 (.033) | .046 (.010) | .063 (.013) |

Note: Standard error in parenthesis.

**Table 3.** Summary statistics for ERP analyses. Standard error in parenthesis.

|  | FRN Peak Latency |  | FRN Mean Amplitude |  |
| --- | --- | --- | --- | --- |
|  | *Win* | *Lose* | *Win* | *Lose* |
| *Win > Lose* | 278 (5) | 276 (6) | 0.24 (0.41) | -0.23 (0.33) |
| *Win = Lose* | 277 (6) | 269 (5) | 0.32 (0.41) | -0.25 (0.35) |
| *Win < Lose* | 277 (6) | 274 (5) | 0.32 (0.37) | -0.19 (0.30) |

|  | FRN Peak Latency |  | FRN Mean Amplitude |  | P3a Peak Latency |  | P3a Mean Amplitude |  |
| --- | --- | --- | --- | --- | --- | --- | --- | --- |
|  | *Actual Win* | *Actual Lose* | *Actual Win* | *Actual Lose* | *Actual Win* | *Actual Lose* | *Actual Win* | *Actual Lose* |
| *Expect Win* | 267 (5) | 271 (5) | 0.39 (0.35) | -0.05 (0.28) | 336 (9) | 357 (8) | 0.84 (0.32) | 0.56 (0.25) |
| *Expect Lose* | 276 (6) | 264 (5) | 0.24 (0.45) | -0.12 (0.32) | 336 (9) | 339 (9) | 0.99 (0.43) | 0.69 (0.40) |

**Figure 2.**
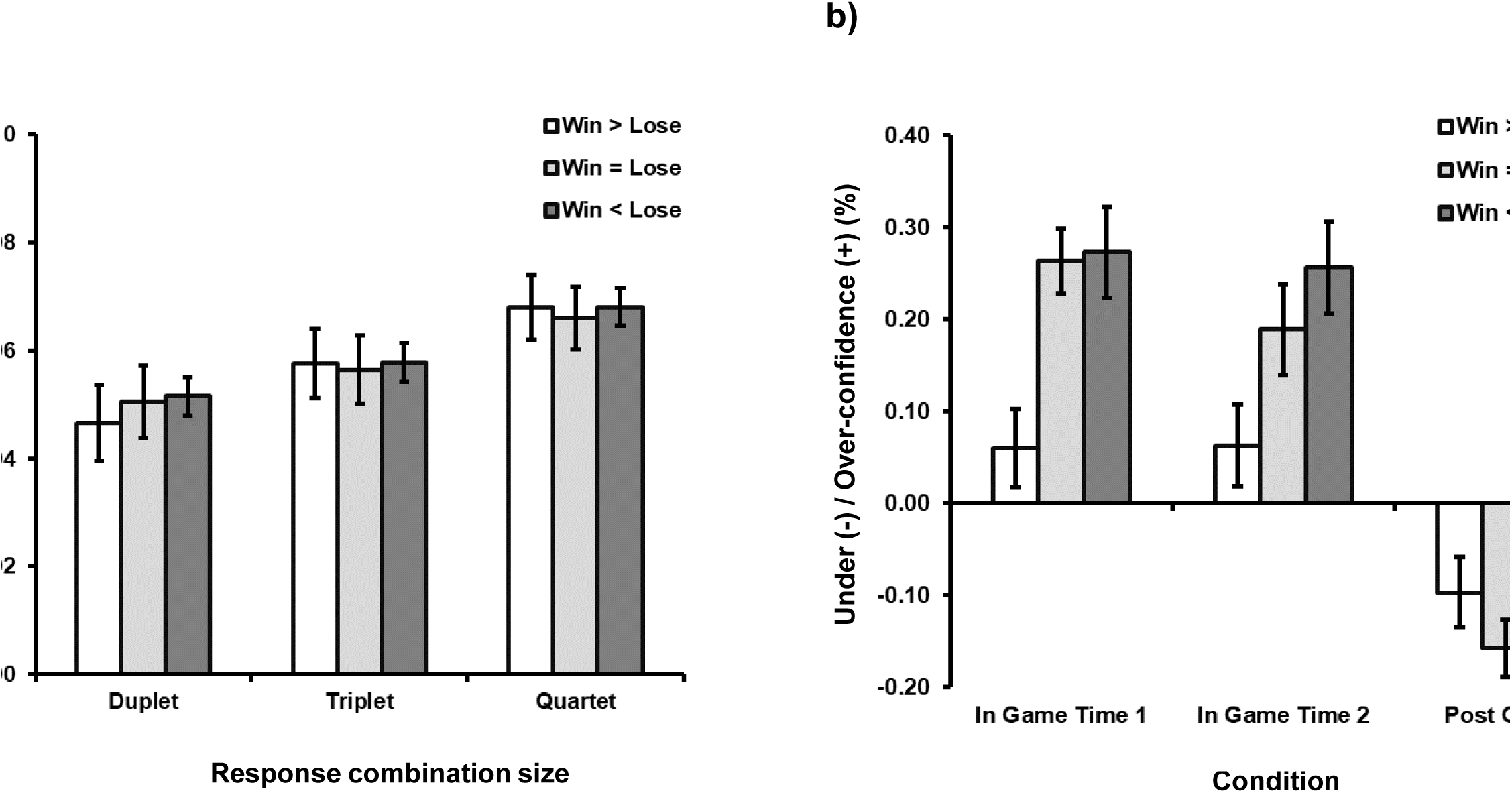
Behavioural data showing a) combinatorial biases as deviations from randomness as a function of response combination size (duplet, triplet, quartet) and weighting of wins and losses in the three conditions (*win > lose, win = lose, win < lose*), and, b) expressions of under- or over-estimate of win rate relative to actual win rate relative to different time points during (in-game time 1, in-game time 2) and after (post game) each of the three conditions. Error bars represent standard error.

Finally, under- or over-confidence as a function of on-line or off-line judgments and win rate was explored by comparing the difference between the actual and predicted win rate at two points during (*in-game time 1, in-game time 2*), and, one point after (*post-game time*) each of the three conditions (*win > lose, win = lose, win < lose*; see Table 1 and Figure 2b). There were main effects of condition: F(2,60) = 9.97, MSE =.044, *p* <.001, □_p_^2^ =.250, and time point: F(2,60) = 49.21, MSE =.061, *p* <.001, □_p_^2^ =.621, as well as an interaction between condition x time point: F(4,120) = 6.41, MSE =.028, *p* <.001, □_p_^2^ =.176. As confirmed by Tukey’s HSD test’s (*p* < .05), participants were generally more over-confident in the *win < lose* (+15.02%) and *win = lose* (+9.94%) conditions, relative to the *win > lose* (+1.47%) condition. Furthermore, participants were equally over-confidence at *in-game time 1* (+20.61%) and *time 2* (+17.65%), which was significantly different from the under-confidence expressed during post-game measurement (−11.83%). The interaction appeared to arise from the slight reduction in over-confidence at *in-game time 2* during the *win = lose* condition.

### ERP Data

FRN peak amplitude (average across all conditions 275 ms; see Figure 3a) showed no significant main effect of condition: F(2,60) = 0.71, MSE = 385, *p* =.497, □_p_^2^ =.023, outcome: F(1,30) = 1.91, MSE = 478, *p* =.178, □_p_^2^ =.060, nor interaction : F(2,60) = 0.64, MSE = 355, *p* =.529, □_p_^2^ =.210. FRN mean amplitude showed reliably greater negativity for negative outcomes (loses; −0.22 uV) relative to positive outcomes (wins; 0.30 uV): F(1,30) = 13.09, MSE = 0.95, *p* =.001, □_p_^2^ =.304, in the absence of a main effect of condition: F(2,60) = 0.03, MSE = 1.61, *p* =.968, □_p_ ^2^ = .001, and in the absence of an interaction : F(2,60) = 0.05, MSE = 0.57, *p* =.948, □_p_^2^ =.002.

**Figure 3.**
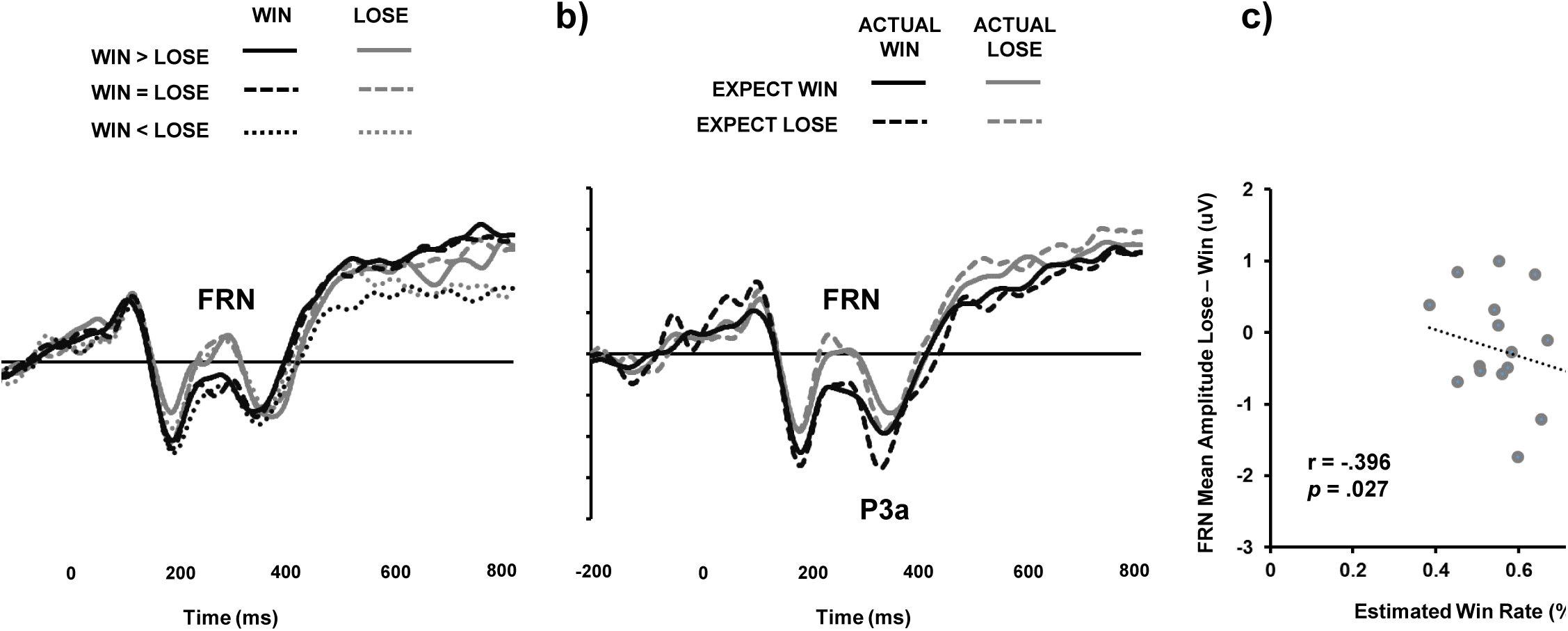
Group-average ERP from 9 fronto-central sites generated by a) the presentation of trial outcome (win, lose) according to weighting of wins and losses in the three conditions (*win > lose, win = lose, win < lose*), and, b) the interaction between expected (*win, lose*) and actual (*win, lose*) trial outcome. In both cases, 20 Hz filter is applied for presentation. c) Correlations between estimated win-rate and FRN mean amplitude difference between loses and wins averaged across three conditions.

FRN was also analysed according to the relationship between the expected (win, lose) and actual (win, lose) outcome of each trial (see Figure 3b). Due to the inability to control the distribution of observations within these four cells, only 24 participants were available for analysis, with observations additionally collapsed across all three conditions ^2^ . FRN peak latency (average 270 ms) showed no significant main effect of expected outcome: F(1,23) = 0.11, MSE = 287, *p* =.745, □_p_^2^ =.005, actual outcome: F(1,23) = 1.10, MSE = 358, *p* =.305, □p^2^ =.046, nor interaction between expected x actual outcome: F(1,23) = 3.98, MSE = 270, *p* =.058, □_p_^2^ =.148. FRN mean amplitude continued to show greater negativity for actual negative outcomes (loses; −0.22 uV) relative to actual positive outcomes (wins; 0.30 uV) in the smaller sample: F(1,23) = 4.94, MSE = 1.18, *p* =.036, □_p_ ^2^ =.177. There was no main effect of expected outcome: F(1,23) = 0.01, MSE = 0.79, *p* =.907, □_p_^2^ <.001, nor an interaction : F(1,23) < 0.01, MSE = 0.45, *p* =.953, □_p_^2^ <.001. P3a peak latency showed a main effect of actual outcome (F(1,23) = 5.39, MSE = 751, p =.030, _p_^2^ =.190), and interaction between actual x expected outcome (F(1,23) = 5.98, MSE = 367, *p* =.023, □_p_^2^ =.206), in the absence of a main effect of expected outcome (F(1,23) = 2.50, MSE = 571, *p* =.128, □_p_ ^2^ =.097). Here, the peak for *expect win – actual loss* (356 ms) was delayed relative to all other cells: *expect win – actual win* (334 ms), *expect loss – actual win* (335 ms), and, *expect loss – actual loss* (339 ms; Tukey’s HSD, *p* <.05). P3a mean amplitude failed to produce any significant effects: actual outcome (F(1,23) = 3.68, MSE = 0.71, *p* =.068, □_p_^2^ =.138), expected outcome (F(1,23) = 0.90, MSE = 1.71, *p* =.351, □_p_ ^2^ =.038), actual x expected outcome (F(1,23) = 0.26, MSE = 0.47, *p* =.614, □_p_ ^2^ =.614). Finally, larger estimated win-rates from all responses across all conditions were correlated with overall larger differences in FRN amplitude between losses and wins (*r* = −.396, *p* =.027; see Figure 3c). Individual correlations within each condition did not provide any further reliable information regarding the source of this effect (*win > lose, r* = −.343, *p* =.059; *win = lose, r* = −.239, *p* =.196; *win < lose, r* = −.321, *p* =.078).

## Discussion

To examine the behavioural and neural interactions between objective and subjective performance during competitive decision-making, participants completed a Matching Pennies game where win-rates were fixed within three conditions (win > *lose, win* = *lose, win* < *lose*). We tested the idea that higher-quality decision-making is more likely when the individual advances from a previous position of success rather than failure. By fixing win-rates across conditions, we exactly controlled for objective performance and hence individual exposure to wins and losses. The quality of decision-making within these contexts was established by comparing optimal, mixed-equilibrium strategy (*random*) behaviour against actual performance where participants potentially deviated from randomness in three ways via item, contingency and combination biases. Overall tendencies to exhibit predictability were shown for all three measures. The item bias was for heads, thereby replicating the reachability account (primary effect) put forward by Bar-Hillel et al. (2014). Contingency biases were revealed for both *win-stay* / *lose-shift* behaviour, thereby replicating the central tenants of operant conditioning. However, *lose-shift* behaviour was not more likely than *win-stay* behaviour, contradicting the findings of Dyson et al. (2016), and, Forder & Dyson (2016). The robustness of both repetition following success and change following failure are currently attributed to the use of two, clearly delineated positive and negative outcomes in MP (i.e., *win* versus *lose*), relative to the use of three outcomes in RPS biased towards negative rather than positive outcomes (i.e., *lose, draw* versus *win*). A more direct comparison between these two paradigms is currently underway to provide direct support for this proposal (Sundvall & Dyson, in preparation).

In contrast to item and contingency biases, combination biases also showed an interaction between the degree of randomness deviation and condition, where participants were closer to expressing randomness with respect to response pairs (i.e., HH, HT, TH, TT) within a condition where wins were more likely than losses. We take this as further evidence that higher-quality decision-making is more likely when exposed to a context containing more positive than negative outcomes (Dyson et al., 2016; Laakasuo et al., 2015; Mitzenmacher & Upfal, 2005). We also observed a second effect where increased exposure to wins lead to more accurate performance. In the case of estimating the likelihood of winning or losing, participants in the *win > lose* condition provided a subjective estimate of performance much closer to actual objective performance (+1.47%). This was in contrast to both the *win = lose* and *win < lose* conditions, where participants exhibited large degrees of over-confidence (+9.94% and +15.02%, respectively). It is worth noting again that these differences in optimism were expressed during in-game periods where all win-rates were 50%. Collectively, these data represent an interesting distinction between attempts to maintain System 2 operation due to accurate positive expectations and unrealistic optimism, respectively (after Shepperd, Pogge & Howell, 2017). In the case of *win > lose*, participants objectively encountered more wins than loses and so did not call upon the secondary mechanism of an *optimism bias* (Cazé & van der Meer, 2013; Sharot et al., 2007) in an attempt to self-regulate current performance. However, in the *win = lose* and *win < lose* conditions, unrealistic optimism was required in order to redress the balance of the objectively lower win-rates encountered in these conditions. In both cases however, the state of belief regarding the experience of more positive than negative outcomes was met. Both of these routes appear to achieve the same goal in attempting to maintain the less predictable, higher-quality output of System 2 over and above the more predictable, lower-quality output of System 1. Our final observation with respect to the behavioural data is that optimism biases were only in evidence during the game and not after the game (see Figure 2b). We take this data as further support that optimism biases are an attempt to self-regulate processing quality *at the time of current performance*. The observation that more sober estimates of success are provided after game performance for all conditions suggests that the charged nature of competition is reduced in ‘post-match analysis,’ along with a similarly reduced need for self-regulation. Other explanations regarding the difference between in-game and post-game estimates stem from the use of multiple, binary decisions in-game and a one-shot, frequency estimate post-game, for which, further study is required.

Our FRN data showed that the initial neural registration of outcome operates in a purely binary fashion, where ‘worse-than-expected’ outcomes (i.e., *loses*) exhibit larger negativity than ‘better-than-expected’ outcomes (i.e., *wins*; e.g., Dyson et al., in review; Forder & Dyson, 2016; Gentsch, Ullsperger & Ullsperger, 2009; Holroyd et al., 2006; Kreussel, Hewig, Krestchmer, Hecht, Coles, & Miltner, 2012; Kujawa, Smith, Luhmann & Hajack, 2013; Nieuwenhuis, Holyord, Mol & Coles, 2004). In our study, FRN did not modulate according to the frequency of losses (Cohen, Elger, & Ranganath, 2007; Forder & Dyson, 2016), nor did it reliably register matches and mismatches between expected and actual outcomes (Holyrod et al., 2003; Oliveria et al., 2007). There was no evidence that such distinctions between expected and actual performance were registered by later ERP components such as P3a (Hajack et al., 2005), and an examination of anterior to posterior mid-line ERP also provided little indication that P3b modulation was exhibited at parietal sites (see Supplementary Graph A). This naturally raises questions regarding our experimental design, and whether the manipulation of win-rate was too small to demonstrate reliable effects. Indeed, the 64% - 36% split employed in the current study was smaller than the 75% - 25% or 80% - 20% splits previously reported (Cohen et al., 2007; Fielding et al., 2018; Hajack et al., 2005). In defence of our choice of parameter, we have four observations. First, the 18% difference between positive and negative outcome was sufficient to observe behavioural differences. Second, we had previously observed modulation of *lose* FRN amplitude across conditions where the number of losses averaged around 33.3% but the value of loses changed (Forder & Dyson, 2016). Therefore, it is not necessary for the number of wins and losses to change in order to observe FRN modulation. Third, previous studies have demonstrated an effect of FRN amplitude at a single trial level, where larger amplitudes for negative outcomes are reported when those outcomes go on to predict behavioural change as opposed to repetition (Cohen & Ranganath, 2007). Fourth, if FRN modulation operates at both a phasic and tonic level, then neural signatures that only distinguish between large-scale differences (in the magnitude of 25% - 30%) would be of little computational value.

Our final empirical observation relates to the correlation between the overall degree of optimism expressed by our participants during the study, and, the distinction between loses and wins exhibited by individual FRN mean amplitude. This correlation once again supports the contention that FRN operates in accordance with a ‘worse-than-expected’ versus ‘better-than-expected’ decision-rule (Holroyd et al., 2006; Kujawa, Smith, Luhmann & Hajack, 2013). For individuals who exhibit high win-rate expectation (larger differences between *win* and *lose* estimates), the differences between positive and negative outcomes should be magnified as shown by the larger difference in FRN mean amplitude for *losses* – *wins*. In contrast, for individuals who exhibit lower win-rate expectation (smaller differences between *win* and *lose* estimates), the distinction between positive and negative outcomes should be less salient, and this is represented by smaller difference in FRN mean amplitude for *losses* – *wins.* As such, the data suggest that if an individual exhibits larger self-belief in success, this will also be manifest in larger neural distinctions between positive and negative outcomes.

An accurate sense of ownership regarding the successes and failures we experience is essential for the maintenance of self-worth and well-being, but this goal is set against a backdrop of cognitive biases that can give rise to various illusions of control (Miller & Ross, 1975; Robins & Beer, 2001; Weary, 1978). In the current context, participants were unable to control the win rates they experienced in any meaningful way and yet still exhibited differences in performance belief across conditions. If optimism biases operate in conditions where individuals have no control, this suggests a central role for performance belief in allocating resources to internal systems that ultimately determine the quality of decision-making and behaviour. Moreover, studying the attribution of wins and losses to internal and external factors is important as it extends beyond the remit of cognitive neuroscience, experimental psychology or economics. In the context of education, students might devalue their self-worth as a result of negative assessment feedback rather than use the information to improve their skills (Crocker & Knight, 2005). In the context of industry, the setting of performance-based rather than learning-based goals (Dweck & Leggett, 1988) may produce inefficient solutions and unethical behaviour as a result of the fear of forthcoming failure (Seijts & Latham, 2005). In the context of problem gambling, the negativity we experience following failure can be greater than the positivity following success even when the objective value of wins and losses are equivalent (Thaler, Tversky, Kahneman & Schwartz, 1997), giving rise to sub-optimal decision-making and potentially catastrophic performance following loss (e.g., Laakasuo et al., 2015; Mitzenmacher & Upfal, 2005). We must understand the mechanisms that give rise to the knowledge that our losses were due to our lack of ability (and know to do better) as well as when our wins were due to luck (and know not to make too much of it), and the current study represents a step towards that larger goal.

## Supporting information

Reproducible Data

## Acknowledgements

Correspondence should be addressed to: Ben Dyson, Department Of Psychology P-217 Biological Sciences Building, University of Alberta, Edmonton, AB, T6G 2E9, Canada.

## Footnotes

1 Individual duplet, triplet and quartet responses had different expected values: 25% for duplets (4 possible responses), 12.50% for triplets (8 possible responses), and, 6.25% for quartets (16 possible responses). This resulted in a natural inflation of the average, absolute deviation from randomness for smaller response types. For example, if an individual simply responded heads for all trials, the average, absolute value was .375 for duplets, .219 for triplets, and, .117 for quartets. To avoid the confound of naturally smaller estimates of deviation for larger response groups, the raw score for duplets was multiplied by.5834 and the raw score for quartets was divided by .5834. Thus, in our extreme hypothetical example where an individual consistently responses heads, this resulted in average, absolute values of .219 for duplets, .219 for triplets, and, .219 for quartets.

2 4 participants could not be included in the analysis due to at least one missing cell. 3 further participants were omitted from analysis if either the ratio of expected win-actual win / expected loss-actual win, or, expected win-actual loss / expected loss-actual loss exceeded 10:1.

