## Supplementary A for "Behavioural and neural interactions between objective and subjective performance in a Matching Pennies game"

### Slide 1
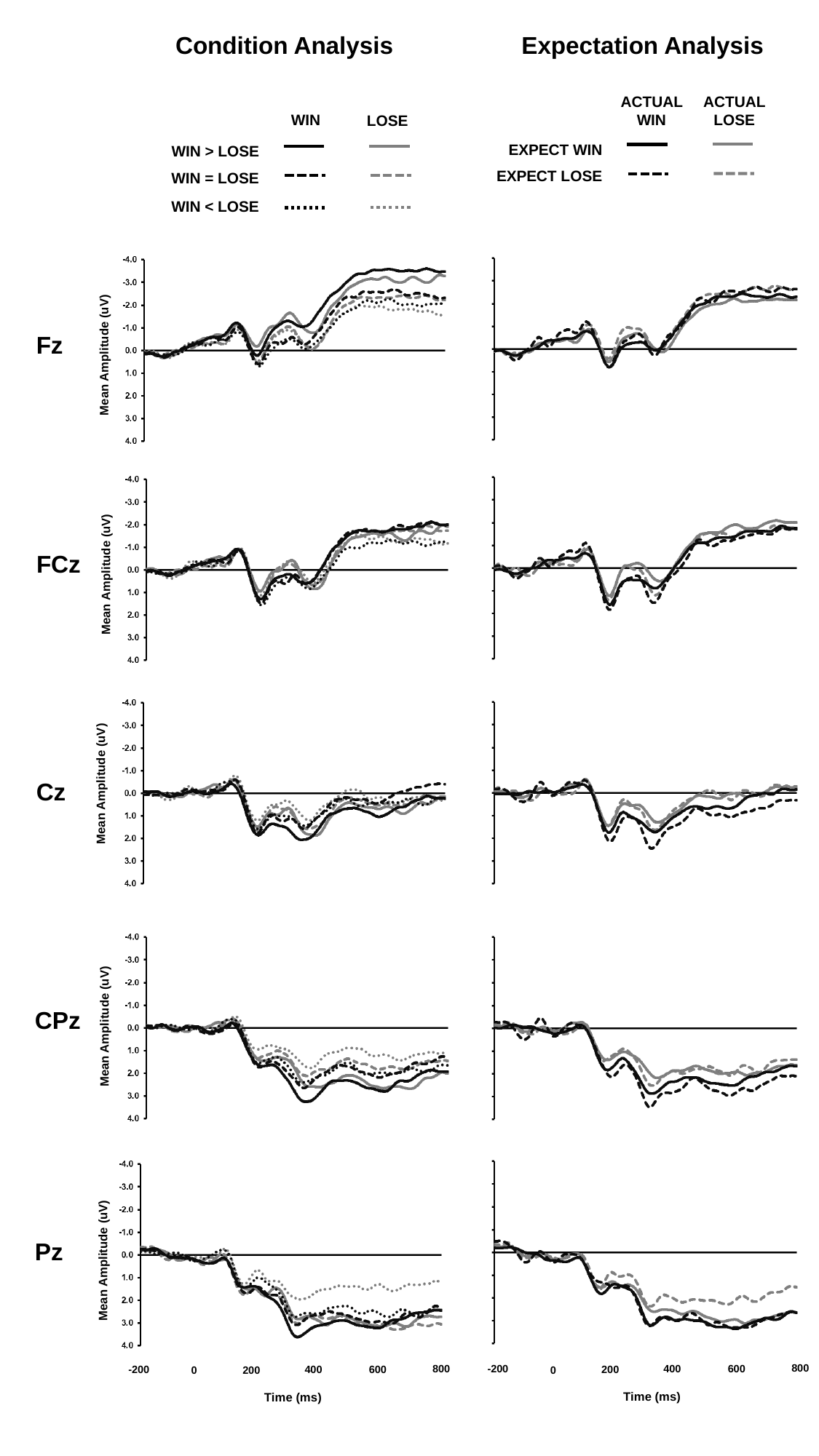

Condition Analysis
Expectation Analysis
ACTUAL WIN
ACTUAL LOSE
WIN
LOSE
EXPECT WIN
WIN > LOSE
EXPECT LOSE
WIN = LOSE
WIN < LOSE
Fz
Mean Amplitude (uV)
FCz
Mean Amplitude (uV)
Cz
Mean Amplitude (uV)
CPz
Mean Amplitude (uV)
Pz
Mean Amplitude (uV)
800
-200
400
800
600
-200
400
200
600
0
200
0
Time (ms)
Time (ms)
